# Cingulate cortex facilitates auditory perception under challenging listening conditions

**DOI:** 10.1101/2023.11.10.566668

**Authors:** Kelsey L. Anbuhl, Marielisa Diez Castro, Nikki A. Lee, Vivian S. Lee, Dan H. Sanes

## Abstract

We often exert greater cognitive resources (i.e., listening effort) to understand speech under challenging acoustic conditions. This mechanism can be overwhelmed in those with hearing loss, resulting in cognitive fatigue in adults, and potentially impeding language acquisition in children. However, the neural mechanisms that support listening effort are uncertain. Evidence from human studies suggest that the cingulate cortex is engaged under difficult listening conditions, and may exert top-down modulation of the auditory cortex (AC). Here, we asked whether the gerbil cingulate cortex (Cg) sends anatomical projections to the AC that facilitate perceptual performance. To model challenging listening conditions, we used a sound discrimination task in which stimulus parameters were presented in either ‘Easy’ or ‘Hard’ blocks (i.e., long or short stimulus duration, respectively). Gerbils achieved statistically identical psychometric performance in Easy and Hard blocks. Anatomical tracing experiments revealed a strong, descending projection from layer 2/3 of the Cg1 subregion of the cingulate cortex to superficial and deep layers of primary and dorsal AC. To determine whether Cg improves task performance under challenging conditions, we bilaterally infused muscimol to inactivate Cg1, and found that psychometric thresholds were degraded for only Hard blocks. To test whether the Cg-to-AC projection facilitates task performance, we chemogenetically inactivated these inputs and found that performance was only degraded during Hard blocks. Taken together, the results reveal a descending cortical pathway that facilitates perceptual performance during challenging listening conditions.

**Significance Statement:** Sensory perception often occurs under challenging conditions, such a noisy background or dim environment, yet stimulus sensitivity can remain unaffected. One hypothesis is that cognitive resources are recruited to the task, thereby facilitating perceptual performance. Here, we identify a top-down cortical circuit, from cingulate to auditory cortex in the gerbils, that supports auditory perceptual performance under challenging listening conditions. This pathway is a plausible circuit that supports effortful listening, and may be degraded by hearing loss.

## Main text / Introduction

Sensory perception often occurs under challenging conditions, such a noisy acoustic background or dim visual environment, yet stimulus sensitivity can remain unaffected. One plausible explanation is that additional cognitive resources are engaged under challenging conditions. For auditory perceptual tasks, including speech comprehension, this concept is referred to as *listening effort*. However, the central mechanisms that are recruited during listening effort remain uncertain.

Measures of human brain activity (i.e., fMRI, EEG/MEG) during challenging listening tasks suggest candidate brain regions that may facilitate performance. For instance, one fMRI study tracked the brain activity of subjects listening to vocoded speech in which the spectral information ranged from intact (i.e., clear) to degraded (Erb et al. 2013). The ‘easy’ listening condition, using clear speech stimuli, activated the AC only. In contrast, difficult listening conditions that contained degraded stimuli, recruited cingulate cortex and other regions associated with executive function. Furthermore, similar effects were observed for a non-speech task. In fact, many fMRI studies report that the cingulate cortex is recruited during performance under challenging listening conditions, including speech comprehension (Eckert et al. 2009), speech with degraded spectral content (Wild et al. 2012), speech intelligibility during multi-talker babble (Vaden et al. 2013), and phoneme discrimination (Gennari et al. 2018). Finally, combined fMRI and electrophysiological measurements suggest that the cingulate cortex modulates the AC while listeners perform auditory tasks (Crottaz-Herbette and Menon 2006).

Perceptual deficits associated with sensory dysfunction are often attributed to degraded processing along the pathway from sensory periphery to primary sensory cortex axis (D. H. and T. N. Wiesel Hubel, n.d.; Van der Loos and Woolsey 1973; Knudsen, Knudsen, and Esterly 1984; Hensch 2005; de Villers-Sidani et al. 2007; Popescu and Polley 2010; Cheetham and Belluscio 2014; Mowery, Kotak, and Sanes 2015). However, a separate literature has emphasized that cognitive mechanisms (e.g., attention, working memory, decision making) can also be disrupted following sensory dysfunction (Moore, Zobay, and Ferguson 2020; Reichman and Healey 1983; Pisoni and Cleary 2003; Holmes, Kitterick, and Summerfield 2017; Dupuis et al. 2015; Hood and Atkinson 1990). For example, a common complaint from individuals with hearing loss is that extra *listening effort* is required to understand speech even when normal audibility is restored (Pichora-Fuller et al. 2016; Sarampalis et al. 2009; Hornsby 2013; Fred H. Bess Benjamin W. Y. Hornsby 2014). While moderate effort is required for optimal performance in normal hearing conditions (McGinley, David, and McCormick 2015), persistently high effort due to hearing loss can contribute to communication errors and give rise to elevated mental fatigue, anxiety, social withdrawal, and depression (Morata et al. 2005; Alhanbali et al. 2018).

The rodent cingulate cortex is also associated with effortful tasks. For example, inactivating rat Cg resulted in reduced willingness to expend greater effort to obtain a high-value reward (Hart et al. 2017, 2020; Holec, Pirot, and Euston 2014; Hosking, Cocker, and Winstanley 2014). Recordings of Cg neural activity also reveal selective responses to behavioral conditions that require specific effort-based demands (Hart et al. 2020; Porter, Hillman, and Bilkey 2019). *These observations suggest that the cingulate cortex is a cortical region that may be recruited during task performance under effortful listening conditions.* Here, we asked whether the gerbil cingulate cortex (Cg) sends a projection to the AC, and whether it mediates listening effort on an auditory effort-based behavioral task. Our experiments reveal a strong descending projection from the cingulate to the auditory cortex. Local inactivation of cingulate cortex degraded performance on an auditory perceptual task, but only for trials in which task difficulty was high. Furthermore, selective inactivation of cingulate inputs to auditory cortex also degraded performance when task difficulty was high. Overall, our results provide causal evidence that the cingulate cortex plays a role in maintaining sensory perceptual skill in the face of challenging sensory decisions.

## Results

### Animals achieve comparable thresholds for easy and hard blocks on a listening effort behavioral task

To assess listening effort, gerbils were trained to perform an amplitude modulation (AM) rate discrimination task (see Methods). Using an appetitive Go-Nogo paradigm, adult gerbils discriminated between “Go” stimuli (AM rates from 4.5-12 Hz, broadband noise carrier, 100% depth) and a “Nogo” AM stimulus (4 Hz; Figure 1A). Trials were clustered into ‘Easy’ or ‘Hard’ blocks in which the sound duration was 1s or 0.25s, respectively (Figure 1B). For each animal, two psychometric functions were generated within a testing session: one for Easy blocks and one for Hard blocks. Figure 1C shows psychometric functions collected from all animals (n=19). AM rate discrimination thresholds were determined from psychometric functions, where the AM rate discrimination threshold was defined as the value at which the signal detection metric, *d’*, was equal to 1. For all animals, the average (± standard error) AM rate threshold for Easy blocks was 4.9 ± 0.03 Hz and the average threshold for Hard blocks was 5.1 ± 0.05 Hz (Figure 1D; one-way repeated measures ANOVA; *F(1,36)*=1.08, p=0.31). False alarm (FA) rates also did not differ significantly between Easy and Hard blocks (Easy=21.5; Hard=15.7; one-way repeated measures ANOVA, *F(1,36)* = 3.97, p=0.06). However, there was a significant effect of stimulus block on psychometric function slope, where Hard blocks yielded shallower slopes (Figure 1E; Easy: 0.35 ± 0.004 *d’*/Hz; Hard: 0.19 ± 0.005 *d’*/Hz; one-way repeated measures ANOVA, *F(1,36)*=27.4, p=0.00006). We interpret these findings as indicating that animals are less certain during Hard blocks, but increased listening effort permits them to obtain the same sensitivity as during Easy blocks. If so, then interference with a neural mechanism that supports listening effort should selectively diminish performance during the hard blocks, as tested below.

**Figure 1.**
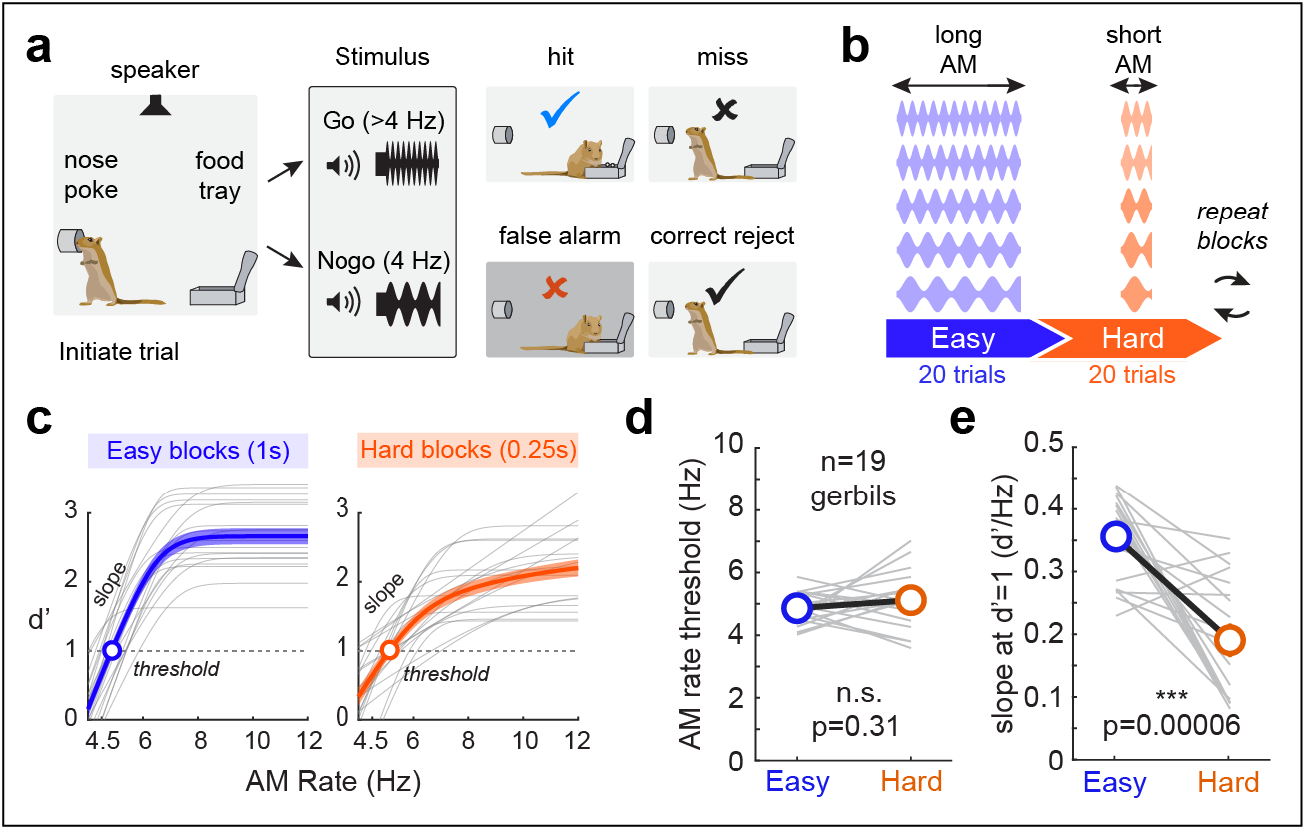
Behavioral task to assess auditory effort. (**a**) Schematic of amplitude modulation (AM) rate discrimination task used to probe auditory effort. Using an appetitive Go-Nogo paradigm, gerbils were trained to discriminate between “Go” stimuli (4.5-12 Hz AM rates; broadband noise carrier, 100% modulation depth) and a “Nogo” stimulus (4 Hz AM). (**b**) Stimulus parameters (AM rate, sound duration) were varied to adjust the difficulty of listening conditions. Trials were clustered into easy (blue) or hard (orange) blocks, where the sound duration was 1s or 0.25s, respectively. (**c**) Psychometric functions for easy (top) and hard (bottom) blocks for a group of animals (n=19). Gray lines indicate individual functions and thick lines are the group average (± standard error). We define threshold as the AM rate at which the signal detection metric, *d’*, is = 1 (circle). The slope of the function is drawn from the slope (*d’*/Hz) at threshold (*d’*=1). (**d**) AM rate thresholds (n=19) are not different between blocks (left; p=0.27, one-way repeated measures ANOVA), whereas the slope of the functions are shallower with hard blocks (right; p<0.001, one-way repeated measures ANOVA).

### Cingulate cortex projects to auditory cortex

Since the literature suggests a role for cingulate cortex during effortful decisions (see Introduction), we first asked whether it sends a robust anatomical projection to the gerbil AC. Retrograde and anterograde virus tracers were injected into AC and cingulate cortex, respectively. First, retrograde vector viruses (AAVrg-hSyn-mCherry) were injected into AC at superficial (300 μm) and deep (800 μm) locations below the cortical surface (Figure 2A,B). Cross-sections through the cingulate cortex reveal retrogradely-labeled cell bodies in the subregion 1 of the cingulate cortex (Cg1; Figure 2C), and were restricted to Layers 2/3 (Figure 2D). Retrograde viruses were also injected into other auditory cortical regions including the anterior auditory field (AAF; Figure 2E,F). Here, AAVrg-hSyn-mCherry was injected into primary AC (A1), and a different retrograde vector was injected into AAF (AAVrg-hSyn-EGFP). Retrograde labeling from primary AC (A1)- and AAF-recipient neurons were present in Cg1 cell bodies (Figure 2G) which were restricted to Layers 2/3 (Figure 2F). Finally, a retrograde vector (AAVrg-hSyn-EGFP) was injected into the dorsal auditory cortex (AuD; Figure 2I,J) and demonstrated a direct projection to Cg1 neurons across all cortical layers (Figure 2K,L). To confirm these findings, we injected an anterograde vector (AAV1-hSyn-EGFP) into Cg1 (Figure 2M,N) and identified afferent labeling within AAF, AC, and AuD, both superficial (∼200-300 μm) and deep (∼800 μm) layers (Figure 2M,O,P). Taken together, these experiments revealed a strong, descending projection from layer 2/3 of the Cg1 to superficial and deep layers of dorsal and primary AC.

**Figure 2.**
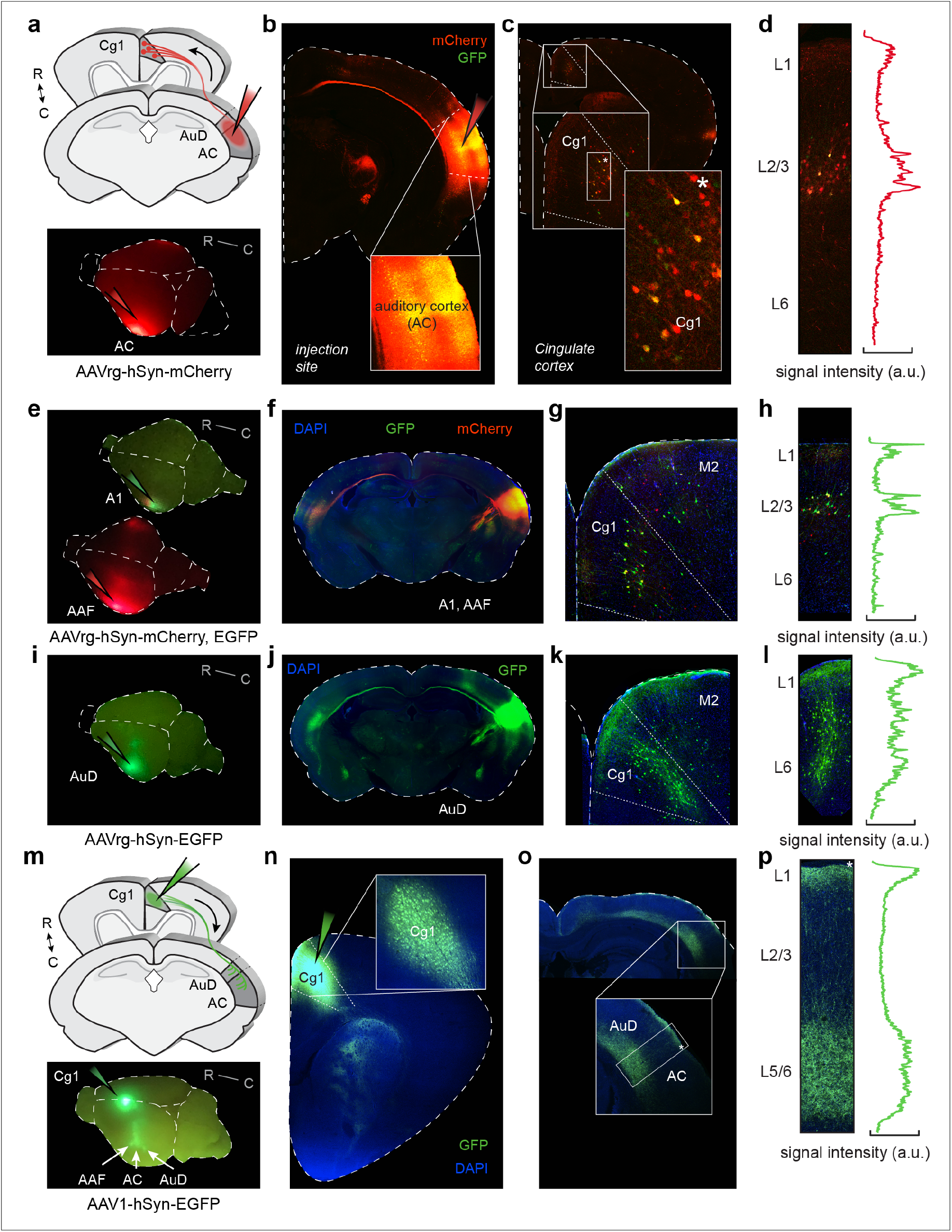
Cingulate cortex neurons project to auditory cortex. (**a**) Schematic showing retrograde AAVrg-hSyn-mCherry injection into auditory cortex (AC). Inset: Whole-brain view of injection site. (**b**) Cross-section through injection site in AC. (**c**) Cross-section through cingulate cortex (Cg). Retrograde labeling from AC-recipient neurons are present in cell bodies of Cg subregion 1 (Cg1). (**d**) Labeled Cg1 cell bodies are restricted to layers 2/3. (**e**) Whole-brain view of retrograde injection sites into primary AC (A1; AAVrg-hSyn-EGFP) and the anterior auditory field (AAF; AAVrg-hSyn-mCherry). (**f**) Cross-section through injection site in A1, AAF. (**g**) Cross-section through Cg1. Retrograde labeling from A1- and AAF-recipient neurons are present in Cg1 cell bodies. (**h**) Labeled Cg1 cell bodies are restricted to layers 2/3. (**i**) Whole-brain view of retrograde injection sites into dorsal AC (AuD; AAVrg-hSyn-EGFP). (**j**) Cross-section through injection site in AuD. (**k**) Cross-section through Cg1. Retrograde labeling from AuD-recipient neurons are present in Cg1 cell bodies. (**l**) Labeled Cg1 cell bodies are present in layers 2/3 through layer 6. (**m**) Schematic showing anterograde AAV1-hSyn-EGFP injection into Cg1. Inset: Whole-brain view of Cg1 injection site. Projections to AuD, AC, AAF are visible (see arrows). (**n**) Cross-section through injection site in Cg1. (**o**) Cross-section through AC where anterograde labeling from Cg1 is present. (**p**) Anterograde labeling is restricted to superficial (L1) and deep (L5/6) layers of AC.

### Local inactivation of cingulate cortex diminishes performance only during hard stimulus blocks

We next determined whether the cingulate cortex contributes to task performance on the AM discrimination task during Hard blocks. To do so, we reversibly silenced cingulate cortex with bilateral infusion of muscimol, a selective gamma-aminobutyric class A receptor (GABA_A_R) agonist. Since muscimol at high concentrations can impede motor function (Laviola and Alleva 1990), we titrated the dose needed for the cingulate cortex inactivation experiments (Supplementary Figure 1) and identified the optimal dose (0.0344 mg/mL).

Trained gerbils (n=9) were implanted with bilateral cannulae above Cg1. After a 1 week recovery, baseline performance was re-established (”No drug” sessions). On successive days, direct infusions of muscimol or saline were delivered 30 minutes prior to behavioral testing on the listening effort task (see timeline in Figure 3A). Each animal was tested across 1-3 sessions for muscimol infusions and 1-3 sessions of saline infusions. After the final session, anterograde virus (AAV1-hSyn-EGFP) was infused to confirm cannulae locations within Cg1 (Fig 3B). Figure 3C shows the average psychometric functions for Easy blocks (1s sound duration) and Figure 3D shows the average functions for Hard blocks (0.25s sound duration) for the three experimental conditions: No Drug (black), Muscimol (magenta), and Saline (blue). The thin lines indicate individual functions, and the thick lines show the group average (shaded area: standard error). Following local inactivation of cingulate cortex, animals displayed AM rate discrimination thresholds for Easy blocks that were comparable to No Drug and Saline control conditions (Figure 3E,F). The average thresholds were: No Drug, 4.8 ± 0.11 Hz AM; Muscimol: 4.96 ± 0.19 Hz AM; and Saline: 5.01 ± 0.18 Hz AM. A two-way mixed model ANOVA indicates no main effect of infusion on performance during Easy blocks (*F(2,39)* = 0.64, p=0.53). However, for Hard blocks, psychometric performance was disrupted following cingulate inactivation. Figure 3G shows the average AM rate threshold for each individual infusion day (No Drug, Muscimol 1, Saline 1, and so forth), and Figure 3H shows the average threshold for each condition. On average, thresholds were elevated following muscimol infusion as compared to Saline infusion. The average thresholds were: No Drug, 5.2 ± 0.35 Hz AM; Muscimol: 6.2 ± 0.5 Hz AM; and Saline: 5.2 ± 0.3 Hz AM. A two-way mixed model ANOVA revealed a main effect of infusion group on performance during Hard blocks (*F(2,37)* = 4.14, p=0.02), followed by a post-hoc test for multiple comparisons (Tukey HSD) which found Muscimol thresholds were significantly elevated compared to Saline controls (*t*= 2.61, p=0.03), whereas Saline and No Drug did not differ significantly (*t*= 0.02, p=0.99).

**Figure 3.**
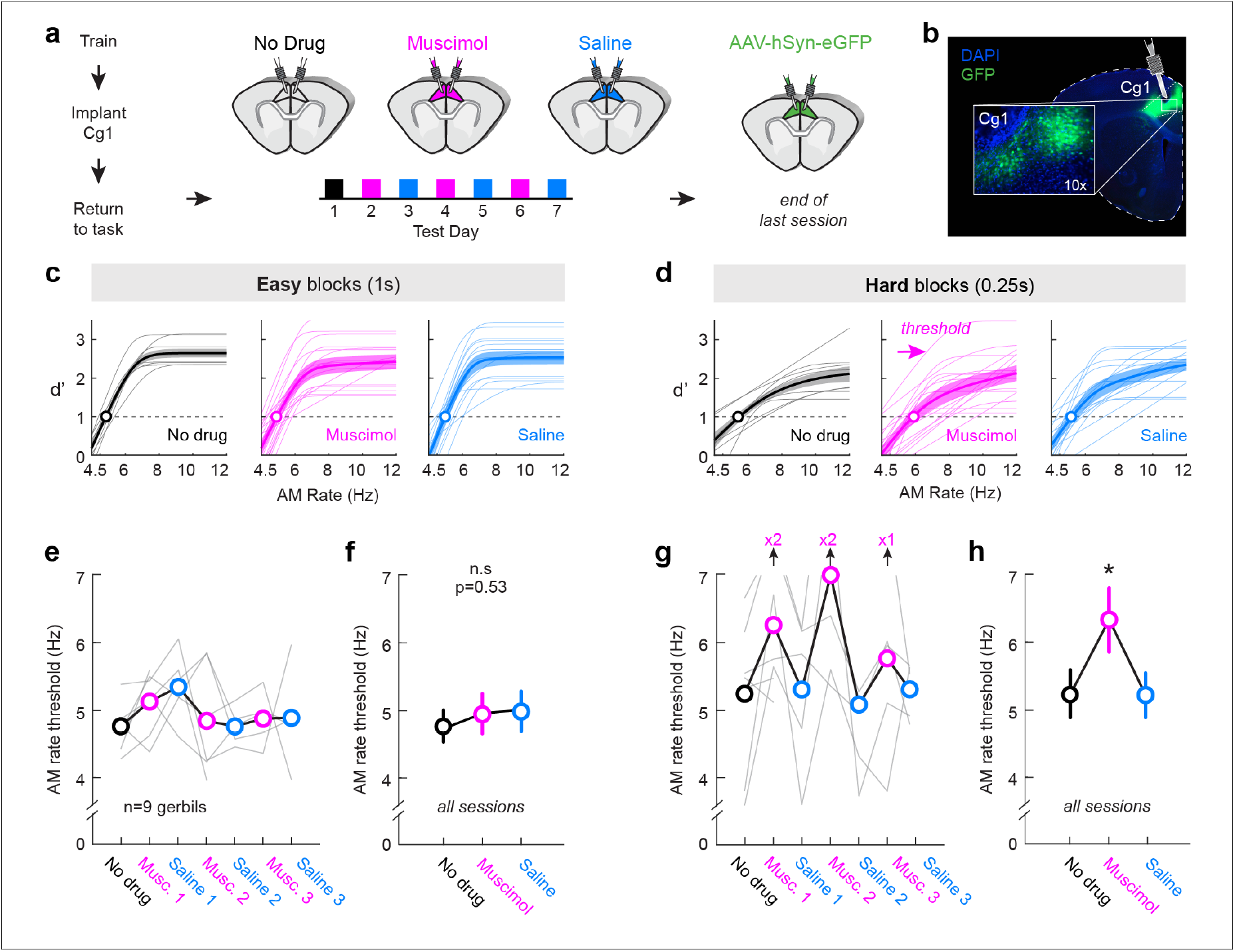
Local inactivation of Cingulate cortex disrupts performance only during hard stimulus blocks. (**a**) Timeline of experimental procedures. Behaviorally-trained gerbils (n=9) were implanted with bilateral cannulae in Cg1. After recovery, baseline performance was re-established (”No drug”, black). On successive days, muscimol (magenta) or saline (blue) infusions were delivered prior to testing on auditory effort task. (**b**) After the final session, GFP virus (AAV-hSyn-eGFP) was infused to confirm pharmacological target location in Cg1. (**c, d**) Average psychometric functions across all animals (thick lines) for each experimental condition (ND, Muscimol, Saline) for Easy blocks (c) and Hard blocks (d). Shaded regions indicate the group average ± standard error (SE). (**e**) AM rate threshold for (Hz) across all testing days for Easy blocks. Circles indicate the group average and gray lines indicate individual data. (**f**) Average AM rate threshold (± SE) for all animals and testing sessions combined (Easy blocks only). Thresholds across infusion conditions for Easy blocks are not statistically different from one another (Two-way mixed model ANOVA, p=0.53). (**g**) AM rate threshold across all testing days for Hard blocks. Sessions with elevated thresholds greater than 7 Hz AM are indicated with an upwards arrow. (**h**) Average AM rate threshold (± SE) for all animals and testing sessions combined (Hard blocks only). A two-way mixed model ANOVA revealed a main effect of infusion group (p=0.02). A post-hoc test for multiple comparisons revealed thresholds for Hard Muscimol blocks are significantly elevated compared to Saline controls (p=0.03).

Elevated discrimination thresholds for Hard blocks could be attributed to elevated false alarm (FA) rates and/or lower hit rates. During easy blocks FA rates remain low during Easy blocks regardless of infusion condition (No Drug: 12.9 ± 2.6; Muscimol: 23.4 ± 4.7; Saline: 16.8 ± 3.9; mixed model ANOVA, *F(5, 78)* = 2.54, p=0.03; followed by Tukey HSD test for multiple comparisons: p=0.53-0.99). For Hard blocks, FA rates are elevated following Muscimol infusions (No Drug: 21.8 ± 4.1; Muscimol: 30.6 ± 7; Saline: 18.0 ± 5.2; FA data not shown). A two-way mixed model ANOVA found a main effect of infusion condition (*F(5, 78)* = 2.54, p=0.03), though a post hoc test for multiple comparisons found no significant difference between conditions (Muscimol-No Drug: *t*= 1.4, p=0.7; Muscimol-Saline: *t*= 2.5, p=0.13; No Drug-Saline: *t*= 0.6, p=0.99). For Hit rates, inactivating cingulate cortex does not alter rates for Easy blocks (mixed model ANOVA followed by Tukey HSD test for multiple comparisons: p=0.07-0.99). For Hard blocks, there is a significant main effect of infusion condition (*F(5, 412)* = 4.4, p=0.0006), though a post hoc test for multiple comparisons found no significant difference between conditions (Muscimol-No Drug: *t*= 1.2, p=0.86; Muscimol-Saline: *t*= 1.7, p=0.5; No Drug-Saline: *t*= 0.23, p=0.99; Hit rate data not shown). Since inactivating cingulate cortex yielded modest changes to FA and Hit rates for Hard blocks, we next examined the signal detection variable, criterion (zHit+zFA)/2), which quantifies the behavioral response regardless of trial type (ie., Go, Nogo). Positive criterion values indicate a more liberal response strategy (reported “yes” regardless of trial type) and negative values indicate a more conservative response strategy (reported “no” regardless of trial type). We found that inactivating cingulate cortex shifted response types for Easy blocks (No Drug: −0.1 ± 0.08; Muscimol: +0.12 ± 0.1; Saline: −0.08 ± 0.1) and Hard blocks (No Drug: −0.14 ± 0.1; Muscimol: +0.1 ± 0.2; Saline: −0.2 ± 0.1) to a more liberal strategy (mixed model ANOVA: *F(5, 484)* = 6.6, p<0.0001; post hoc test for multiple comparisons: MuscimolEasy-SalineEasy: p=0.02, MuscimolHard-SalineHard: p=0.0008; MuscimolHard-NoDrugHard: p=0.03). Though for Easy blocks, there was not a significant difference between Muscimol and No Drug criterion values (p=0.08; criterion data not shown).

To rule out potential confounds we first asked whether muscimol infusion led to a decrease in trial number. Neither the number of Go trials (Supplementary Figure 2A), nor Nogo trials (Supplementary Figure 2B) differed during Easy or Hard blocks. For Go trials, a mixed model ANOVA followed by a test for multiple comparisons found no difference between Easy Muscimol and control conditions (Muscimol-NoDrug, p=0.06; Muscimol-Saline: p=0.7) or between Hard Muscimol and Saline (p=0.6). For Nogo trials, a mixed model ANOVA followed by a post hoc test for multiple comparisons indicated no differences between conditions for Easy (p=0.23-0.99) or Hard (p=0.4-0.99) blocks. Thus, Go and Nogo trial numbers cannot account for the performance deficits following cingulate cortex inactivation. We then asked whether task performance was influenced by the isoflurane anesthesia used ≥30 minutes prior to testing. Therefore, we took a group of trained animals (n=5) and exposed them to the concentration and timing of a typical isoflurane exposure followed by a 30-minute recovery period. Animals then performed the Auditory effort Task, and exhibited perceptual thresholds similar to what they achieved following No Drug and Saline infusion sessions (mixed model ANOVA: *F(5, 30)* = 1.4, p=0.27; Supplementary Figure 3A,B).

Finally, we asked whether bilateral cingulate cortex inactivation was required to diminish performance during Hard blocks. Unilateral cannulae were implanted above one Cg1 in a group of behaviorally-trained gerbils (n=4). Following recovery, baseline performance was re-established (No drug), and muscimol or saline infusions were delivered unilaterally prior to testing (2-7 sessions for each animal; Supplementary Figure 4A). We found that thresholds for Easy and Hard blocks remained unaffected following unilateral muscimol infusions in Cg1 compared to No Drug and Saline controls (mixed model ANOVA, *F(5, 32)* = 0.94, p=0.47; Supplementary Figure 4B). Thus, both cingulate cortex regions need to be inactivated to give rise to performance deficits on the Auditory effort Task.

### Selective Inactivation of Auditory cortex-projecting Cingulate neurons diminishes performance only during hard stimulus blocks

Although local inactivation of cingulate cortex resulted in poorer performance during hard stimulus blocks on the listening effort task, it is likely that many downstream targets were impaired. To test whether the cingulate projection to AC is required for psychometric performance on Hard blocks, we chemogenetically inactivated AC-projecting cingulate neurons. Trained gerbils (n=8) received bilateral AC (A1 and AuD) injections of a retrograde adenovirus that transfects pyramidal neurons with HM4Di, an inhibitory designer receptors exclusively activated by designer drug (DREADD) receptor, and bilateral cannulae were implanted over Cg1. Baseline performance was re-established (No Drug) over the subsequent 3-5 weeks during which optimal virus expression occurred. Prior to each behavioral testing session, direct infusion of Compound 21 (C21; chemogenetic actuator of hM4D) or saline (control) were delivered via cannulae on alternate days (Figure 4A). Each animal completed a total of 6 alternating sessions of C21 and saline infusions (3 sessions each). On the final testing day, no infusions were delivered (No Drug, post infusions). Once behavioral testing was complete, we confirmed the location of the virus injection sites in both auditory cortices (AC, AuD), the bilateral cannulae above Cg1, and the presence of retrogradely-labeled cell bodies in Cg1 (Supplemental Figure 5A). Indeed, the target locations were confirmed in all animals tested (n=8). Supplemental Figure 5B shows an example whole-brain view of the virus injection sites in left and right AC. We verified the presence of HM4Di-mCherry infected AC neurons (Supplemental Figure 5C,D) and established that the cannulae were in the correct location above Cg1 (Supplementary Figure 5E). We also confirmed cell body labeling of retrograde mCherry virus in Cg1 (Supplemental Figure 5F,G).

**Figure 4.**
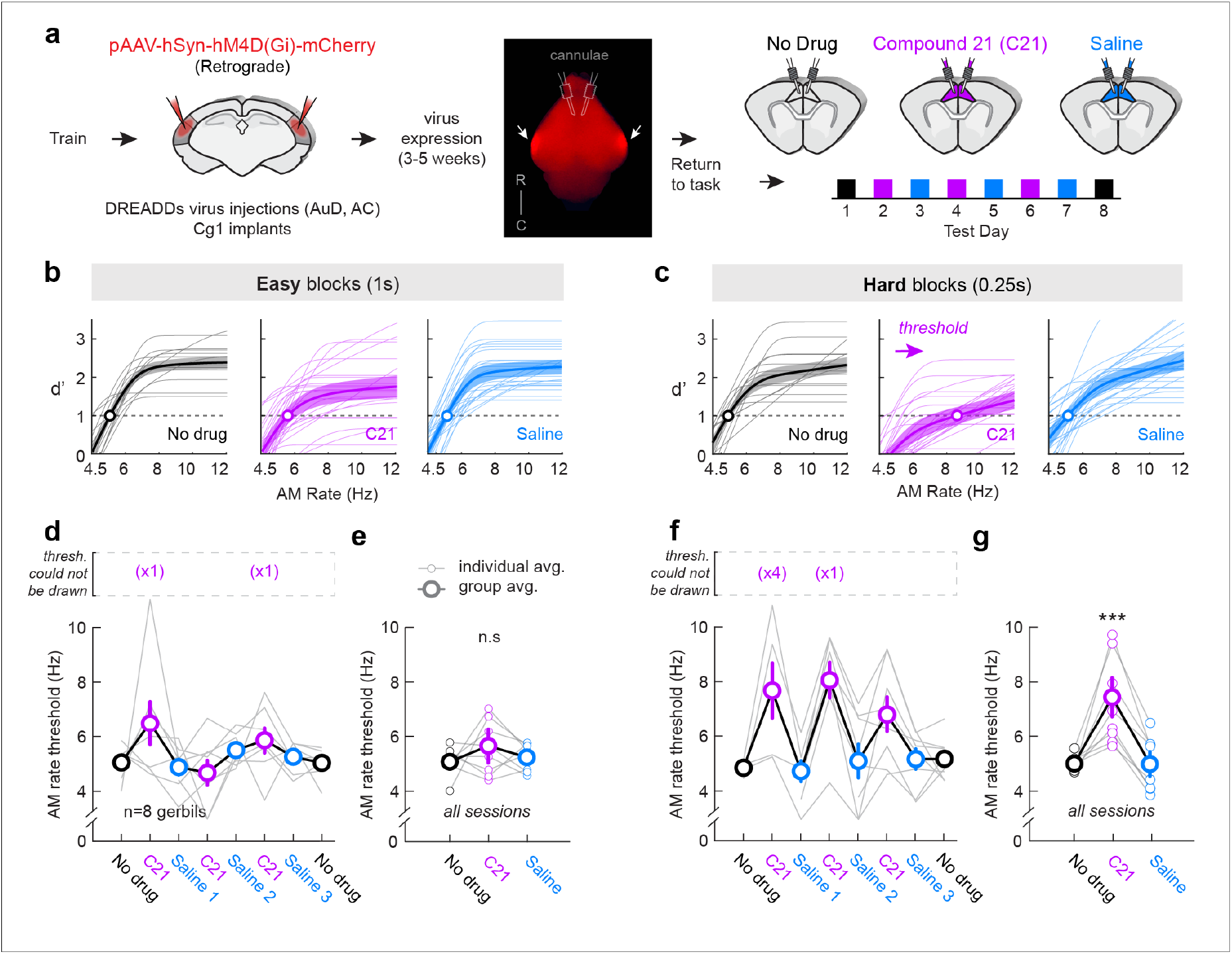
Inactivating Cingulate-to-Auditory cortex-specific projections disrupts performance during hard stimulus blocks. (**a**) Timeline of experimental procedures. Behaviorally-trained gerbils (n=8) received bilateral injections of pAAV-hSyn-hM4D(Gi)-mCherry (retrograde) into auditory cortex (AC and AuD), followed by bilateral cannulae implants in Cg1. During a 3-5 week period (for optimal virus expression; see arrows in whole-brain example), baseline performance was re-established (”No drug”, black), and infusions of Compound 21 (C21, purple) or saline (blue) were delivered prior to behavioral testing sessions. On the final testing day, no infusions were delivered (”No Drug”-post infusions). (**b**) Average psychometric functions across all animals (thick lines) for each experimental condition (No Drug, C21, Saline) for Easy blocks (b) and Hard blocks (c). Shaded regions indicate the group average ± standard error (SE). (**d**) AM rate threshold for (Hz) across all testing days for Easy blocks. Circles indicate the group average and grey lines indicate individual data. Values within the grey dotted rectangle indicate the number of sessions in which threshold could not be drawn (i.e., d’<1). (**e**) AM rate threshold for all animals and testing sessions combined (Easy blocks only). Larger circles depict group average (± SE), and smaller circles indicate individual condition averages. Thresholds across infusion conditions for Easy blocks are not statistically different from one another (two-way mixed model ANOVA followed by a post hoc test for multiple comparisons, p=0.7-0.99). (**f**) AM rate threshold across all testing days for Hard blocks. Circles indicate the group average and grey lines indicate individual data. Values within the grey dotted rectangle indicate the number of sessions in which threshold could not be drawn (i.e., *d’*<1). (**g**) Average AM rate threshold (± SE) for all animals and testing sessions combined (Hard blocks only). A two-way mixed model ANOVA revealed a main effect of infusion condition (p<0.0001). A post-hoc test for multiple comparisons revealed thresholds for Hard C21 blocks were significantly elevated compared to control conditions (p<0.0001).

Psychometric functions for all infusion condition sessions (No Drug, C21, Saline) are shown for Easy and Hard blocks (Figure 4B,C). The thin lines indicate functions for each animal, and the thick lines show the group average (shaded area: standard error). Discrimination thresholds are shown for each individual infusion day (Figure 4D,F) and for all sessions combined (Figure 4E,G). The average discrimination thresholds for Easy blocks were: No Drug, 5.1 ± 0.2 Hz AM; Muscimol: 5.6 ± 0.6 Hz AM; and Saline: 5.2 ± 0.2 Hz AM. For Hard blocks, the average thresholds were: No Drug, 5.02 ± 0.2 Hz AM; Muscimol: 7.4 ± 0.7 Hz AM; and Saline: 5.0 ± 0.4 Hz AM. When comparing thresholds between groups, we found a significant main effect of infusion condition (two-way mixed model ANOVA, *F(5, 115)* = 10.3, p<0.0001). A post-hoc test for multiple comparisons (Tukey HSD) found that animals displayed comparable thresholds for Easy blocks across all infusion groups (C21-No Drug: *t*=1.4, p=0.7; C21-Saline: *t*=1.1, p=0.9; No Drug-Saline: *t*=-0.4, p=0.99; Figure 4E). However, unlike for Easy blocks, inactivating Cg-AC specific projections resulted in a significant elevation of thresholds for Hard blocks compared to No Drug (*t*=5.5, p<0.0001) and Saline (*t*=6.1, p<0.0001) thresholds (Figure 4G).

We next examined the FA and Hit rates to determine whether the elevated thresholds are due to increased FA rates or poorer Hit rates (or both). Supplementary Figure 6A,B shows the average FA rate for Easy and Hard blocks for each infusion condition. The average FA rate for Easy blocks were: No Drug: 19.1 ± 3.1; C21: 28.9 ± 6.0; Saline: 19.6 ± 3.6, and for Hard blocks: No Drug: 22.4 ± 4.6; C21: 42.6 ± 7.4; Saline: 18.1 ± 4.2. A two-way mixed model ANOVA found a significant main effect of infusion group on FA rates (*F(5, 122)* = 9.5, p<0.0001). A post hoc test for multiple comparisons (Tukey HSD) found that Cg-to-AC inactivation had no effect on FA rate for Easy blocks compared to No Drug (*t*=2.1, p=0.3) or Saline control conditions (*t*=2.2, p=0.25) but did yield a significant effect for Hard blocks (C21-No Drug: *t*=4.3, p=0.0006; C21-Saline: *t*=5.8, p<0.0001). For Hit rates, we examined the average Hit rates for all sessions combined for Easy and Hard blocks (Supplementary Figure 6C,D). A two-way mixed model ANOVA found a significant main effect of infusion group on Hit rates (*F(5, 628)* = 4.2, p=0.0009). A post hoc test for multiple comparisons found no significant effect of Cg-to-AC inactivation on Hit rates for Easy blocks compared to No Drug (*t*=-1.5, p=0.7) or Saline controls (*t*=-1.05, p=0.9), but did reveal a significant effect for Hard blocks compared to No Drug Hit rates (*t*=-4.1, p=0.0007), though not compared to Saline Hit rates (*t*=-2.07, p=0.3). The Hit rates for the two control conditions, No Drug and Saline, are not significantly different from one another (*t*=2.2, p=0.2). We also examined the signal detection variable, criterion, to determine whether there were changes following Cg-to-AC inactivation (Supplementary Figure 6E,F). We found a significant main effect of infusion group on the criterion variable (*F(5, 122)* = 2.4, p=0.04). A post hoc test for multiple comparisons identified no effect between groups for Easy blocks (C21-No Drug: *t*=0.6, p=0.99; C21-Saline: *t*=1.01, p=0.92; No Drug-Saline: *t*=0.33, p=0.99) and for Hard blocks, no effect between C21 and No Drug (*t*=1.0, p=0.91) and a significant effect between C21 and Saline (*t*=3.2, p=0.02). The two control conditions, No Drug and Saline, are not significantly different from one another (*t*=1.9, p=0.4). Thus, we attribute the performance deficits observed during Hard stimulus blocks to an elevation of FA rates and poorer Hit rates, with no change to performance strategy (compared to No Drug control).

The elevated thresholds observed following C21 infusions for Hard blocks could be due to differences in Go or Nogo trial number. To rule this out, we plotted the total number of trials for Go trials (Supplementary Figure 7A) and Nogo trials (Supplementary Figure 7B) for each block and for each infusion condition. For Go trials, there were no differences in trial number between infusion conditions for Easy or Hard blocks (*F(5, 122)* = 1.6, p=0.17). For Nogo trials, we did find a main effect of infusion condition on trial number (*F(5, 122)* = 3.4, p=0.006), though a post hoc test for multiple comparisons found that for Easy blocks there was no significant difference between C21 and No Drug (*t*=-0.2, p=1.0) or Saline (*t*=1.7, p=0.54). For Hard blocks, there was no difference between C21 and No Drug (*t*=1.1, p=0.87), though there was a difference between C21 and Saline (*t*=2.9, p=0.05). There were no differences between the two control conditions, No Drug or Saline (*t*=1.5, p=0.66). Thus, differences in Go and Nogo trial numbers cannot account for the performance deficits following Cg-to-AC inactivation.

The gerbil AC spans nearly ∼2mm in the rostral-caudal dimension (Radtke-Schuller et al. 2016). For the DREADDs experiment described in Figure 4, all retrograde virus injections were made in more rostral AC locations (∼2.1 mm re: Lambda). However, it was unclear whether targeting more caudal AC regions would also give rise to performance deficits following Cg1 inactivation with C21. Therefore, we injected retrograde DREADDS adenovirus into caudal AC (+1.4 mm re: Lambda) of two trained animals (Supplementary Figure 8A), and bilateral cannulae were implanted over Cg1. Baseline performance was re-established (No Drug) over the subsequent 3-5 weeks during which optimal virus expression occurred. Prior to each behavioral testing session, direct infusions of Compound 21 (C21; chemogenetic actuator of hM4D) or saline (control) were delivered via cannulae on alternate days, just prior to each behavioral testing session. Each animal completed a total of 6 alternating sessions of C21 and saline infusions (3 sessions each). Supplementary Figure 8B confirmed the target virus location in caudal AC. We also found sparse labeling of retrograde signal in Cg1 neurons (Supplementary Figure 8C). Supplementary Figure 8D shows the AM rate thresholds for all animals and all sessions combined. Following C21 infusions in Cg1, while we found a main effect of infusion condition (*F(5, 28)* = 2.9, p=0.03, a post hoc test for multiple comparisons revealed no effect on thresholds for Easy or Hard blocks compared to No Drug and Saline controls (Easy blocks: C21-No Drug, *t*=0.89, p=0.95; C21-Saline, *t*=-0.14, p=1.0; Hard blocks: C21-No Drug, *t*=1.99, p=0.0.38; C21-Saline, *t*=1.3, p=0.77). We also found no effect of C21 on False alarm rates compared to control conditions (*F(5, 28)* = 0.6, p=0.68; Supplementary Figure 8E). Thus, retrogradely targeting caudal AC does not impair performance on the listening effort task.

Taken together, our results reveal a descending cortical pathway from the cingulate cortex to auditory cortex that, when inactivated, diminishes performance during hard blocks only. Thus, we conclude this cingulate-to-auditory cortex pathway facilitates perceptual performance during listening effort.

## Discussion

Cognitive effort plays a key role in everyday tasks, from making complex decisions to engaging in conversations. While moderate effort is required for optimal performance (McGinley, David, and McCormick 2015; Yerkes and . Dodson 1908), overallocation can lead to cognitive fatigue (Boksem and Tops 2008; Inzlicht, Shenhav, and Olivola 2018). The difficulty of auditory perceptual tasks can vary, depending on the sensory and cognitive processing ability of the listener, acoustic features of the stimulus itself, and environmental challenges (Peelle 2018). Listening effort, defined as the allocation of cognitive resources during auditory tasks (Pichora-Fuller et al., 2016), is thought to improve performance through engagement of mechanisms such as selective attention and working memory. While the neural networks that support listening effort are not well understood, measures of human brain activity (i.e., fMRI, EEG/MEG) suggest that the anterior cingulate cortex is engaged (Erb et al. 2013; Eckert et al. 2009; Wild et al. 2012; Vaden et al. 2013; Gennari et al. 2018; Crottaz-Herbette and Menon 2006). Here, we report that a strong, descending projection from gerbil cingulate to auditory cortex can selectively facilitate perceptual performance during a challenging listening condition.

### Cingulate cortex is recruited during effortful behavioral decisions in humans and rodents

Research on effort-based decision-making has identified a central network, including the anterior cingulate cortex, that is recruited when effort demands are high (Dosenbach et al. 2008; Aben et al. 2020; Umemoto, Inzlicht, and Holroyd 2019). This extends to perceptual tasks in both the visual and auditory domains. For instance, when humans perform a visual detection task with low or high effort demands (i.e., detecting an object with low vs high visual noise interference), the dorsal anterior cingulate cortex is activated during the hard detection trials (Aben et al. 2020). The anterior cingulate cortex is also recruited during challenging listening conditions, including for difficult speech comprehension (Eckert et al., 2009), speech with degraded spectral content (Wild et al., 2012), speech intelligibility during multi-tacker babble (Vaden et al., 2013), and phoneme discrimination (Gennari et al., 2018). One fMRI study tracked the brain activity of subjects listening to vocoded speech in which the spectral information ranged from intact (i.e., clear) to degraded (Erb et al., 2013). The ‘easy’ listening condition, using clear speech stimuli, activated the auditory cortex only. In contrast, difficult listening conditions that contained degraded stimuli, recruited cingulate cortex and other regions associated with executive function. Further, Erb et al. (2013) report that the magnitude of cingulate cortex activity scaled with discrimination difficulty on an amplitude modulation rate discrimination task, the same task used in our experiments. One interpretation of these findings is that the cingulate cortex provides executive support when effort demands are high. If so, then disrupting this region should degrade listening performance. Indeed, we found that reversible inactivation of the gerbil cingulate cortex selectively disrupted perceptual performance during hard stimulus blocks. Gerbils displayed poorer AM rate thresholds for hard blocks (0.25s) following inactivation of cingulate cortex (Figure 3) or selective inactivation of AC-projecting cingulate cortex neurons (Figure 4), implicating a causal role of cingulate cortex on perceptual difficulty (i.e., rate discrimination) and task difficulty (i.e., long vs short AM durations).

A separate literature has identified a role for rodent cingulate cortex (Cg) during effort-based (both physical and cognitive) decision-making tasks. In rats, inactivating Cg resulted in reduced willingness to expend greater physical effort (e.g., climbing a ramp, lever presses) to obtain a high-value reward (Holec et al., 2014; Hart et al., 2017, 2020). In another study, rats performed a visuospatial attention task with low-or high-effort trials to obtain sucrose rewards of varying quantity (Hosking, Cocker, and Winstanley 2014). Similar to our study, they varied the duration of the stimulus to adjust cognitive effort: long (1s) or brief (0.2s) durations of light were used for low-or high-effort trials, respectively. When Cg was inactivated, rats reduced their choice of high-effort trials (i.e., brief duration of light stimulus). This is consistent with our finding that Cg inactivation disrupted performance for hard stimulus trials using short sound stimuli (0.25s). Further, rats performing a visual discrimination task exhibit disrupted decision confidence with more difficult sensory stimuli following Cg inactivation (Stolyarova et al. 2019). Recordings of Cg neural activity also reveal selective responses to behavioral conditions that require specific effort-based demands (Porter et al., 2019; Hart et al., 2020). Overall, our findings are consistent with existing reports in humans and rodents that suggest the Cg is recruited during effortful behavioral decisions.

### Potential mechanisms of cingulate-mediated facilitation of effortful performance

Top-down, descending inputs to sensory cortex can mediate plasticity observed in sensory neurons during behavioral tasks. For example, Cg inputs to mouse primary visual cortex (V1) modulates visual cortex processing during a discrimination task, where optogenetic activation of Cg neurons enhanced V1 neuron responses and improved visual discrimination (Zhang et al. 2014). AC neurons also exhibit changes to response properties based on behavioral context such as with task engagement: Studies often compare AC neuron responses to the same stimuli under two conditions: while the animals attend to the stimulus during task performance (i.e., engaged), or while the animal is passively listening (i.e., disengaged). In one study, AC neurons rapidly change their tuning properties when animals attend to a target tone during a tone-detection task compared to responses when they are passively disengaged (Fritz et al. 2003). In another study, AC neuron responses to a target sound are enhanced when ferrets actively discriminate stimuli on a Go-Nogo auditory task, as compared to passive listening conditions (Bagur et al. 2018). Furthermore, AC neuron responses are enhanced when animals attend selectively to a target sound, whereas responses to distractor stimuli are suppressed (Schwartz and David 2018). Animals that learn positive or negative stimulus reward associations also exhibit rapid, selective suppression or enhancement of AC neuron tuning, respectively, to the target frequency (David, Fritz, and Shamma 2012). Task difficulty can also modulate AC activity: AC neurons become poorly tuned to a target sound when the SNR is more perceptually difficult (Atiani et al. 2009). Rapid changes to AC responses have been proposed to account for improved behavioral performance: when rats are engaged on a Go/Nogo auditory task, AC neurons shift their stimulus-evoked responses and tuning curves, improving auditory detection thresholds (Carcea, Insanally, and Froemke 2017). Similarly, task engagement on an auditory detection task improves detection thresholds of AC neurons and correlates with behavioral performance learning (von Trapp et al. 2016; Caras and Sanes 2017). These observations show that the AC is modulated by behavioral context, and suggest that its processing may be regulated during listening effort. Here, we report a strong, descending projection from L2/3 of Cg1 to superficial and deep layers of primary and dorsal AC. Thus, it is plausible that hard stimulus blocks engage cingulate-mediated modulation of stimulus encoding in the AC to facilitate auditory perception.

### Relationship to anatomical and functional regions of the cingulate cortex

The cingulate cortex in rodents has been divided into structurally and functionally distinct regions, though the nomenclature across reports largely depends on whether the terms used are homologous (e.g., anterior cingulate cortex, ACC; midcingulate cortex, MCC) or nonhomologous (e.g., cingulate area 1, Cg1; cingulate area 2, Cg2; prelimbic cortex, PL; infralimbic cortex, IL) to humans and primates (Francis-Oliveira, Leitzel, and Niwa 2022; van Heukelum et al. 2020; Laubach et al. 2018; Vogt 2016). The ACC and MCC are divided in the rostral-caudal plane and both encompass Cg1 and Cg2, which are divided horizontally in the coronal plane (van Heukelum et al. 2020; Francis-Oliveira, Leitzel, and Niwa 2022). The two subdivisions are associated with functional differences, where ACC is associated with social, emotional (i.e., positive, negative affect) and pain regulation, while MCC is associated with attention and reward-based decision-making (for review, see (van Heukelum et al. 2020). Here, we use the nonhomologous nomenclature typically used for rodents (mice: (Franklin and Paxinos 2008); gerbil: (Radtke-Schuller et al. 2016), as the cingulate-to-auditory cortex specific projections were restricted to Cg1 in gerbils (Figure 2), and did not extend to the entire ACC or MCC regions which include Cg2. Our study suggests that the Cg1 neurons that facilitated performance under effortful listening conditions were primarily at the ACC/MCC border in the gerbil (∼+0.5 mm rostral from bregma).

Although our chemogenetic inactivation of Cg-to-AC projections yielded behavioral deficits, it is possible that the Cg and/or AC are recruited via reciprocal or indirect pathways. For example, Budinger et al., (E. Budinger, Heil, and Scheich 2000; Eike Budinger et al. 2008) report reciprocal connections between gerbil Cg and AC. We also found distinct laminar distribution patterns of Cg inputs to AC which depended on the AC region (primary AC, AuD, AAF; Figure 2). Cg neurons in L2/3, sent projections to superficial (L1) and deep (L5/6) layers of AC and AAF, whereas retrograde injections into dorsal AC yielded labeled Cg neurons across all layers. Neurons in L2/3 output to pyramidal neurons in L5/6, which compute information flowing through the cortical network before sending projections to other cortical and subcortical regions. These include visual cortex (Zhang et al. 2014), thalamus (Zhang et al. 2016), and amygdala (Jhang et al. 2018). Cg also receives inputs from many areas, including other cortical regions (e.g., orbital, medial prefrontal, retrosplenial, parietal), as well as hippocampus, basal forebrain, thalamus, and brainstem (Fillinger et al. 2017). Thus, it is plausible that the behavioral task also engages these indirect networks that involve Cg. Further, other regions may also mediate this top-down influence onto AC, such as frontal cortex (Winkowski et al. 2013) including orbitofrontal cortex (Winkowski et al. 2018; Ying et al. 2023; Mittelstadt and Kanold 2023). In our task, cognitive mechanisms that engaged Cg and other regions noted above, may contribute to aspects of listening effort that could explain our causal findings. This includes arousal, attention-based performance monitoring, perceived difficulty, and auditory working memory (Eckert et al. 2009; Peelle 2018; Dosenbach et al. 2008; de Gee et al. 2022).

### Disentangling effort from attentional and reward-based mechanisms

We have defined listening effort broadly to include all cognitive resources that enhance performance under challenging (i.e., difficult) environmental conditions. In principle, this could include the modulation of selective attention, working memory, arousal, performance monitoring, and others (Poeppel, Mangun, and Gazzaniga 2020). If our manipulation led to auditory inattention alone, then we would expect to see an increase in the number of miss trials and psychometric functions with lower asymptotic values following Cg inactivation, which we did not see (Supplemental Figure 6). A second caveat is that Cg can be modulated by reward (Amiez, Joseph, and Procyk 2006) and willingness to expend physical effort to receive a reward (Holec et al., 2014; Hart et al., 2017, 2020). However, these studies generally have rodents choose between different quantities (i.e., small vs large) or valence of reward (e.g., sucrose vs water). Here, we sought to avoid reward-related biases, and fixed the quantity (2 pellets/trial) and type (food pellet only) of reward for all trials. If we were reducing the value of a reward, then we would expect to see a decrease in the number of initiated Go trials. However, the number of Go trials are the same between control and Cg inactivation conditions (Supplemental Figure 2, 7).

While listening effort is thought to recruit cognitive mechanisms that involve motivation, arousal, and attention, it might be feasible to disentangle these attributes. For instance, the cognitive contributions may be recruited in a distinct, temporal manner across the trial period that allow for the perceptual decision to be made: attention-based mechanisms are generally recruited in anticipation of trial initiation (Buran, von Trapp, and Sanes 2014; Liu, Denton, and Nelson 2007) and during sensory processing (i.e., sound stimulus), followed by a brief post-stimulus processing period during which the animal integrates the sensory information with prior knowledge (i.e., an *effort-specific epoch* influenced by cognitive engagement), and culminates in a behavioral decision (e.g., seek food, re-poke). In support of this idea, a pupil-based proxy for listening effort in humans yields the largest changes in pupil size during the post-stimulus period of a trial: the time window following stimulus presentation and prior to participant response (Winn et al. 2018).

Furthermore, silencing the Cg in rats disrupts the waiting period following a visual stimulus (Stolyarova et al. 2019), suggesting it may be involved in post-stimulus processing of sensory information. Therefore, an important future direction would be to examine cingulate cortex activity across behavioral timescales (i.e., within-trial to block-level) and effort context.

## Materials and Methods

### Experimental subjects

A total of 28 Adult (P≥105) Mongolian gerbils (*Meriones unguiculatus,* 15 females, 13 males) were used in the study. Gerbils were raised from commercially obtained breeding pairs (Charles River Laboratories) and housed on a 12-hour light / 12-hour dark cycle with full access to food and water, unless otherwise noted. All procedures were approved by the Institutional Animal Care and Use Committee at New York University.

### Auditory stimuli

Gerbils were tested on an amplitude modulation (AM) rate discrimination task. The AM stimuli (broadband noise carrier, 100% depth) varied from 4-12 Hz. AM stimulus duration was varied (0.25 or 1s) to modulate task difficulty. AM is a temporal feature present in most natural sounds (Singh and Theunissen 2003) including speech (Shannon et al. 1995).

### Behavioral training and psychometric testing

#### Behavioral apparatus

All behavioral experiments were performed in a sound-attenuating booth (Industrial Acoustics) and were observed through a closed-circuit video monitor (Logitech HD C270 Webcam). Animals were placed in a custom-made, acoustically-transparent plastic test cage positioned underneath a calibrated free-field speaker (DX25TG0504; Vifa) 1 m above the test cage. The testing cage contains a nose port and a food tray (Figure 1A). Food pellets are dispensed via an external pellet dispenser (Med Associates, Inc.). Sound stimuli, food delivery, experimental parameters, and data acquisition are controlled on a PC with custom MATLAB (MathWorks) scripts developed by Dr. Daniel Stolzberg (https://github.com/dstolz/epsych) and using a multifunction processor (RZ6; Tucker-Davis Technologies, TDT).

#### Procedural training

Animals were trained on a positive reinforcement Go-Nogo appetitive conditioning procedure used in the laboratory (von Trapp et al. 2017; Yao and Sanes 2021; Ihlefeld, Chen, and Sanes 2016). Animals were placed on controlled food access, and trained to initiate a trial by poking their nose in a nose port. The first training trials used a “Go” signal (12 Hz AM rate; 5s duration), and a food reward (20 mg pellet) was delivered if gerbils approached the food tray. Once gerbils reliably initiated trials, trials that contain a “Nogo” signal (4 Hz AM) are introduced, and animals learned to avoid the food tray and re-poke to initiate trials. Trials were classified as a Hit (correctly approaching food tray for a Go trial), a Miss (failure to approach food tray for a Go trial), a correct reject (CR; correctly re-poking during a Nogo trial), or a false alarm (FA; incorrectly approaching food tray during a Nogo trial; Figure 1A). Performance was quantified using the signal detection metric, *d’*, defined as: *d’* = z(hit rate) – z(false alarm rate) (Green and Swets 1974). Once gerbils reached a criterion level of performance (FA<30%; d’≥2) for 3 sessions, new Go rates are added during subsequent sessions (10, 8, 6, 4.5 Hz AM).

#### Perceptual testing

Once animals reached the training performance criterion (see above), animals were then perceptually tested on a range of AM rates (4.5, 6, 8, 10, 12 Hz). Psychometric functions were collected by presenting Go trials (4.5-12 Hz), randomly interleaved with Nogo trials (4 Hz). Psychometric thresholds are defined as the AM rate at which *d’*=1. Animals completed perceptual training when they had three sessions that met the criterion level of performance (FA<30%; d’≥2) with all five AM rates.

##### Easy versus Hard blocks

To modulate the auditory effort required to perform psychometric testing, task difficulty was varied using sound duration. Within each testing session, trials were clustered into ‘Easy’ or ‘Hard’ blocks, where the sound duration was 1s or 0.25s, respectively (Figure 1B). The trial and block structure for muscimol experiments were as follows: Each block contained 20 randomized Go and Nogo trials. Blocks alternated between easy and hard stimulus durations, and each session contained ≥ 6 blocks (≥ 3 hard, ≥ 3 easy). For the DREADDs experiments, the trial and block structure were as follows: Each block contained 40 randomized Go and Nogo trials. Each session contained a total of 3 blocks, where Block 1 was easy (1s), Block 2 was hard (0.25s), and Block 3 was easy (1s). Gerbils were tested until asymptotic performance (i.e., stable thresholds) were obtained. Trained animals were either used for further behavioral assessment, or underwent surgical procedures for neural manipulation experiments (i.e., muscimol, DREADDs).

### Behavioral tracking

Behavioral sessions were video recorded. Animal tracking was quantified using SLEAP, a framework for pose tracking via deep learning (Pereira et al. 2022). Networks were trained (1300 manually labeled frames; 13 separate videos), frame predictions were examined, and labels were re-adjusted as needed. The network was re-trained until high accuracy was achieved (<2.8 prediction error on 95% of points).

### Anatomical virus tracing

Anatomical tracers were used to determine projections to and from AC (Yao et al. 2020). Gerbils were anesthetized and secured on a stereotaxic apparatus (Kopf). Retrograde (AAVrg-hSyn-EGFP; AAVrg-hSyn-mCherry; Addgene) and anterograde (AAV1-hSyn-EGFP; Addgene) viruses were used to assess Cg-AC connectivity. Viruses were injected (100-200 nL; 2 nL/sec) into Cg (0.4-0.6 mm rostral; 0.3-0.5 mm lateral to bregma) and/or dorsal AC and/or primary AC (0.9 mm rostral; 4.6-4.8 mm lateral to lambda). For AC regions, two injections were made (in L2/3, L5/6) to determine layer-specific connections. After a 3 week survival, brain tissue was extracted and processed.

### Cannula infusions

When stable thresholds on the behavior task were obtained, bilateral cannulae was implanted in Cg. Gerbils were anesthetized, stereotaxically secured, and one large craniotomy (∼2×4 mm) was made above both Cg target regions and exposing the sagittal sinus in the center of the craniotomy. Great care was taken to avoid puncturing the sagittal sinus during the craniotomy and subsequent removal of the dura. The coordinates for targeting Cg were: +0.5 mm re: Bregma and ±0.87 mm re: midline. We found that Bregma was unreliable as a midline reference, so we opted to use the center of the sagittal sinus as the midline reference. Single guide cannulas (26 gauge; 3mm; PlasticsOne), were implanted above each Cg (oriented at 10°). Dummy cannulae were inserted into each guide to keep area clear of debris. <u>Infusions</u>: Gerbils were lightly anesthetized, and an infusion cannulae (33 gauge; 4mm), connected to PE-50 tubing backfilled with mineral oil, was connected to a glass syringe (10 μL, 1801 Gastight Hamilton; 23-gauge) attached to a programmable pump (NE-1600, New Era). The infusion cannulae were inserted into the guide (depth: 1mm in center of Cg1). Either saline, Muscimol (GABA_A_ receptor agonist), or Compound 21 (C21; chemogenetic actuator of hM4Di) was infused into Cg1 (Figure 3A, 4A).

### Muscimol

First, we titrated the optimal Muscimol dose needed for the cingulate cortex inactivation experiment. Supplementary FIgure 1A shows a top-down view of animal position across a 10-minute Off-task session (no trials) for three different Muscimol doses (0.5, 1, and 2 mg/mL; direct infusions into cingulate cortex). We computed the total distance traveled for each condition, and found that increasing doses of muscimol results in less distance traveled (Supplementary Figure 1C). Further, we wanted to ensure that infusing muscimol did not alter motor function the following day (Supplementary Figure 1B). Indeed, the total distance traveled for sessions 24 hours-pre and post-infusion did not differ (Supplementary Figure 1C). Since the lowest muscimol dose (0.5 mg/mL) used during the Off-task sessions still yielded motor alterations, we significantly reduced the dose to 0.0344 mg/mL which did not impede performance on the task (Supplementary Figure 1D,E). Therefore, we used this dose of Muscimol (0.0344 mg/mL) for cingulate cortex inactivation experiments. Prior to testing, bilateral infusions of Muscimol or Saline were delivered to each Cg1 (0.125 μL to each hemisphere) at an infusion rate of 0.2 μL/minute. After each infusion, we waited 4 minutes prior to removing the infusion cannulae to ensure proper drug delivery to target site. After the infusion, the animals recovered for 30 minutes prior to behavioral testing.

### Designer Receptors Exclusively Activated by Designer Drugs (DREADDs)

DREADDs were used to perturb neural activity of intercortical connections (Yao et al. 2020). Here, DREADDs were used to specifically inactivate Cg neurons that project to AC. Bilateral injections of a retrograde virus (AAVrg-hSyn-hM4D(Gi)-mCherry) were made into AC, which transfect terminal projections with the inhibitory DREADDs receptor, HM4Di. A total of four injections were made into each AC to target all cortical layers and span the rostral-caudal range for AC: Rostral AC injection location: +3.3 mm re: bregma; 300 μL at 0.8 mm depth (targeting layers 5/6) and at 0.3 mm depth (targeting layers 2/3) below cortical surface; Caudal AC injection location: +3.8 mm re: bregma: 300 μL at 0.8 mm depth (targeting layers 5/6) and 0.3 mm depth (targeting layers 2/3) below cortical surface. Following injections, cannula guides were implanted above each Cg using the coordinates described above. After recovery, trained gerbils were infused with either C21 (HelloBio; 5 mg/mL; 0.2 μL to each hemisphere) or saline (0.2 μL to each hemisphere) in each Cg at an infusion rate of 0.2 μL/minute. After each infusion, we waited 4 minutes prior to removing the infusion cannulae to ensure proper drug delivery to target site. After the infusion, the animals recovered for 30 minutes prior to behavioral testing.

### Histology

At the end of each anatomical tracing, Muscimol, or DREADDs experiment, gerbils were given an overdose of sodium pentobarbital (150 mg/kg, intraperitoneal injection) and perfused with 0.01 M phosphate-buffered saline followed by 4% paraformaldehyde. The brains were extracted, post-fixed in 4% paraformaldehyde, and embedded in 6% agar and sectioned at 40-60 µm on a vibratome (Lieca). Tissue sections were mounted on gelatin-subbed glass slides, stained with DAPI (Vectashield Vibrance Antifade Mounting Medium with DAPI, Vector Laboratories). Brightfield and/or fluorescence images were collected (Revolve, Echo; Leica confocal; Olympus BX61VS) to determine fluorophore expression in Cg, AC regions and confirm location of cannulae targeting Cg (Figure 3B, 4B). ImageJ was used to quantify layer-specific expression of fluorophores. For the DREADDs experiment, the mCherry signal was amplified for optimal visualization of signal of virus in the injection site in AC, and cell body labeling in Cg (Supplementary Figure 5). The native mCherry fluorescence was visualized by using the Living Color Red rabbit DS polyclonal antibody (cat #632496) and goat anti rabbit IgG conjugated to Alexa Red 555 (Life Sciences, cat #A21428).

#### Statistics

Statistical analyses and procedures were performed using JMP Pro 16.0 (SAS) or custom-written MATLAB (MathWorks, R2023a) scripts utilizing the Statistics and Curve Fitting Toolboxes. For normally distributed data (assessed using Shapiro-Wilk Goodness of Fit Test), values are given as mean ± SEM. Non-normally distributed data are given as median ± Quartile. Unless otherwise noted, all normally distributed data were assessed using parametric procedures (i.e., ANOVA) followed by appropriate *post hoc* controls for multiple comparisons (Tukey-Kramer HSD) when necessary.

## Abbreviations

AC: auditory cortex
AAF: anterior auditory field
AM: amplitude modulation
Cg: cingulate cortex
AuD: dorsal auditory cortex

## Acknowledgements

We thank Samer Masri for his contributions to the anatomy presented in Figure 2A, and are grateful to other members of the Sanes lab for constructive feedback and support. We also thank Claudia Farb for her help with processing and imaging the anatomy from the DREADDs experiment. Special thanks for Drs. Matthew Winn and Matt McGinley for helpful discussions on earlier versions of this work. This research was supported by NIH 1K99DC020570-01 (KLA) and NIH R01DC020279 (DHS).

## Author Contributions

Designed research and secured funding: KLA and DHS; Behavioral training and testing: KLA, MDC, NAL, and VL; Virus injections (anatomical tracers, DREADDs): KLA, MDC, NAL, VL; Drug infusions: KLA, MDC, NAL; Analysis: KLA; Visualization: KLA and DHS; Writing original draft: KLA and DHS; Review & editing: KLA, MDC, NAL, VL, and DHS.

## Conflict of interest

The authors declare no competing financial interests.

## Supplemental Figures

**Supplementary Figure 1.**
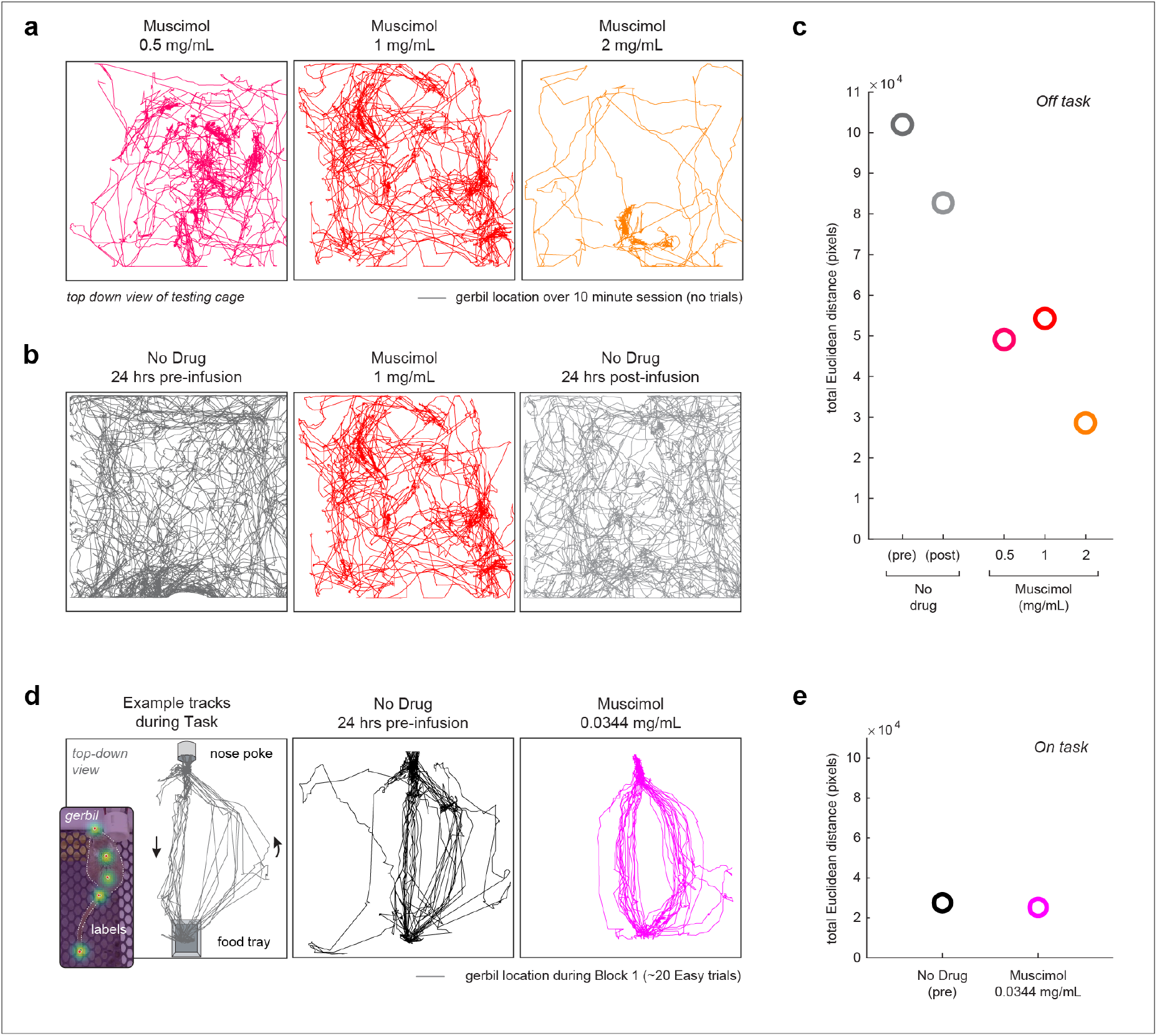
Titrating optimal muscimol dose for cingulate cortex inactivation experiments. (**a**) Top-down view of animal position across a 10-minute Off-task session (no trials) for three different Muscimol doses (0.5, 1, and 2 mg/mL; direct infusions into cingulate cortex). (**b**) Animal tracks for Off-task sessions: prior to infusion (”No Drug, 24 hrs pre-infusion”), following Muscimol infusion (1 mg/mL), and after infusion (No Drug, 24 hrs post-infusion). (**c**) Total Euclidean distance traveled (pixels) for each condition depicted in (a,b). Euclidean distance is calculated using the x- and y-dimensions, where the distance between two points (x1, y1) and (x2, y2) in a Cartesian coordinate system is given by the following formula: Distance = sqrt((x2 - x1)^2 + (y2 - y1)^2). (**d**) Tracks of example animal performing trials during Block 1 of the auditory effort task (”On Task”). Tracks are shown for trials prior to infusions (”No Drug, 24 hrs pre-infusion”) and after infusions using a lower Muscimol dose (0.0344 mg/mL). (**e**) Total Euclidean distance traveled (pixels) for No Drug and Muscimol (0.0344 mg/mL) trials. Both conditions displayed comparable total distance traveled, indicating that the lower Muscimol dose did not alter general motor function. Therefore, this dose of Muscimol (0.0344 mg/mL) was used for cingulate cortex inactivation experiments (Figure 3).

**Supplemental Figure 2.**
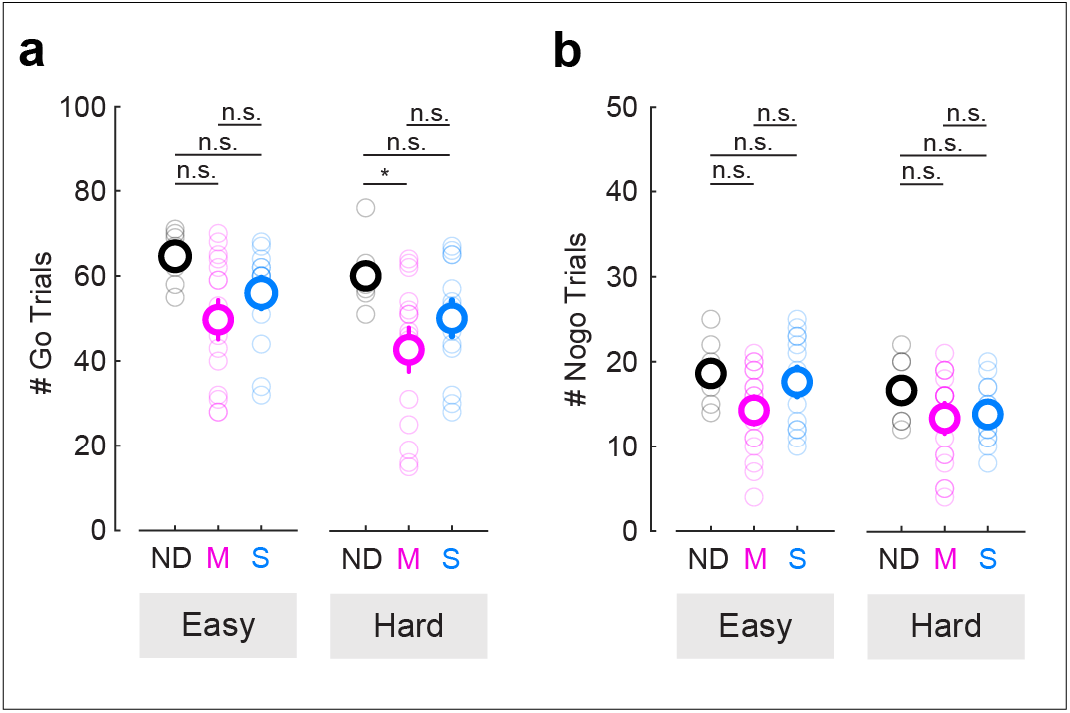
Performance deficits following cingulate inactivation are not due to differences in trial number. (**a**) Number of Go trials for each block (Easy, Hard) for each infusion condition (No Drug, ND; Muscimol, M; Saline, S). Mixed model ANOVA followed by post hoc analysis for multiple comparisons reveal no significant differences between groups for Easy or Hard blocks (p=0.07-0.69), with the exception of No Drug and Muscimol Hard (p=0.02). (**b**) Number of Nogo trials for each block (Easy, Hard) for each infusion condition (No Drug, ND; Muscimol, M; Saline, S). Mixed model ANOVA followed by post hoc analysis for multiple comparisons reveal no significant differences between groups for Easy or Hard blocks (p=0.23-0.99).

**Supplemental Figure 3.**
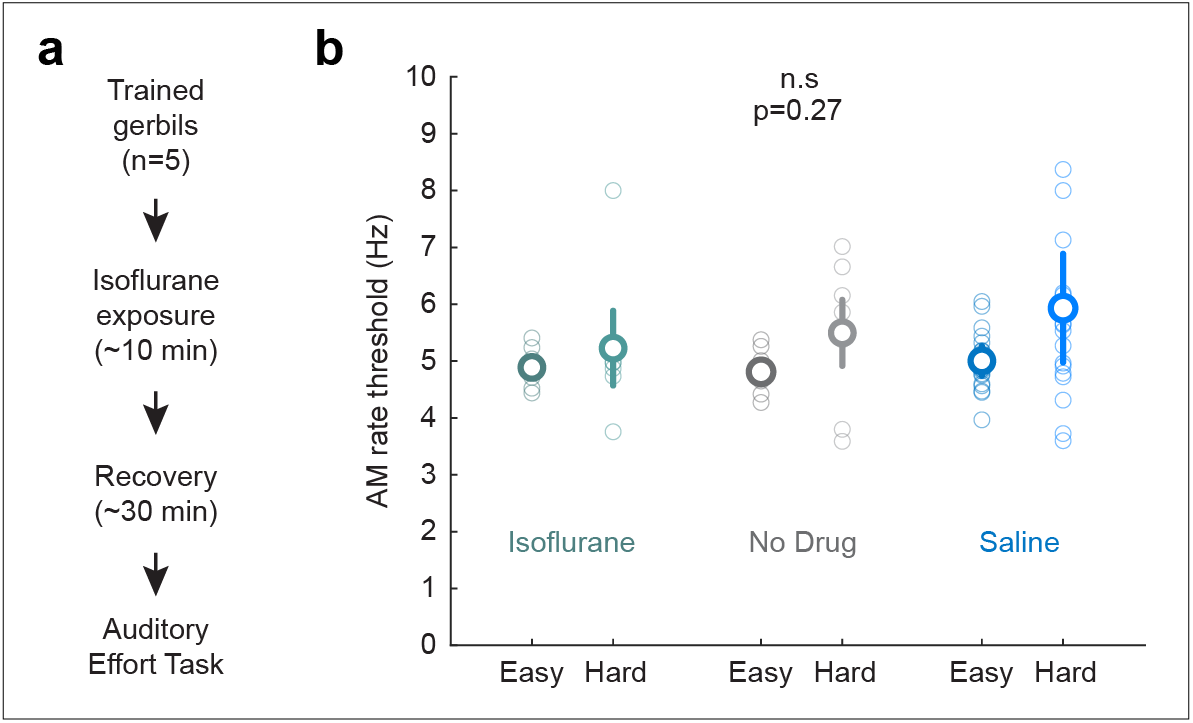
Exposure to Isoflurane does not alter perceptual thresholds. (**a**) Timeline for Isoflurane exposure control experiment. Trained animals (n=5) are exposed to the concentration and timing of a typical isoflurane exposure used in infusion experiments (1-5% Isoflurane, 2% O_2_ over a ∼10 minute span), followed by a 30 minute recovery period. Animals then performed the Auditory Effort Task. (**b**) AM rate threshold (Hz) for Easy and Hard blocks following Isoflurane exposure (n=5). No Drug and Saline infusions thresholds are also shown for the same trained animals (n=5). Isoflurane did not alter AM rate thresholds compared to No Drug and Saline control conditions (mixed model ANOVA, p=0.27).

**Supplemental Figure 4.**
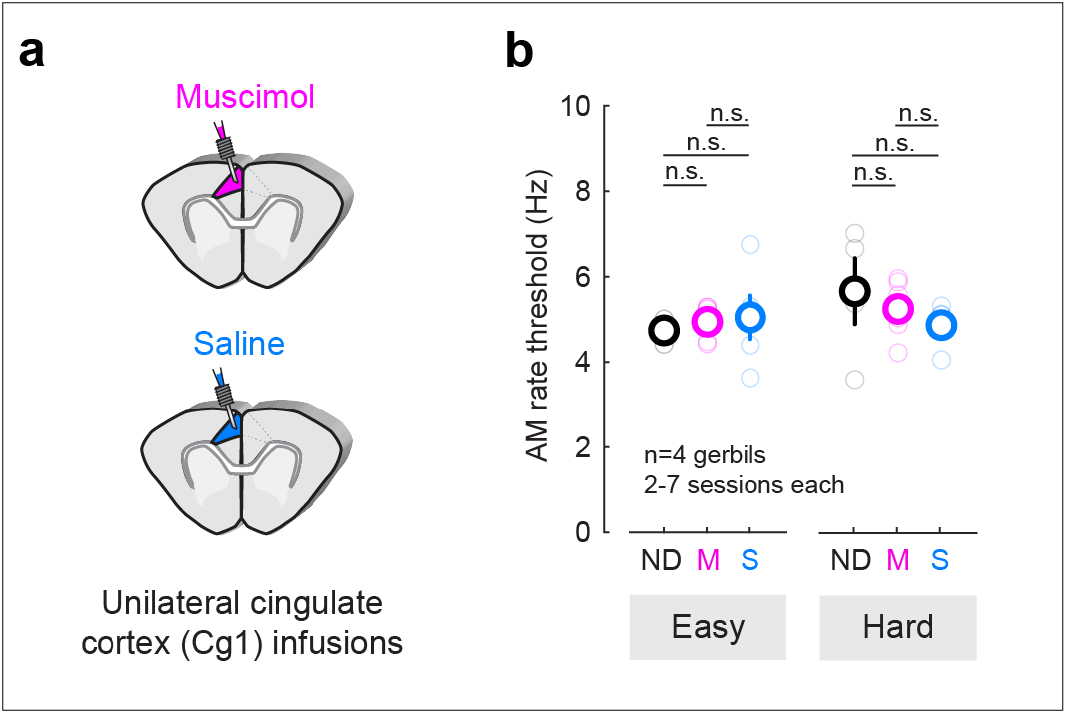
Unilateral cingulate cortex inactivation does not impair performance. (**a**) Schematic of unilateral infusion experiment. Behaviorally-trained gerbils (n=4) were implanted with a unilateral cannulae in one Cg1 side. After recovery, baseline performance was re-established (No drug), and muscimol or saline infusions were delivered unilaterally prior to testing (2-7 sessions for each animal). (**b**) Average AM rate threshold (± SE) for all animals and testing sessions combined. Thresholds remain unaffected following unilateral muscimol infusions in Cg1 compared to No Drug and Saline controls (Mixed model ANOVA, p=0.47).

**Supplemental Figure 5.**
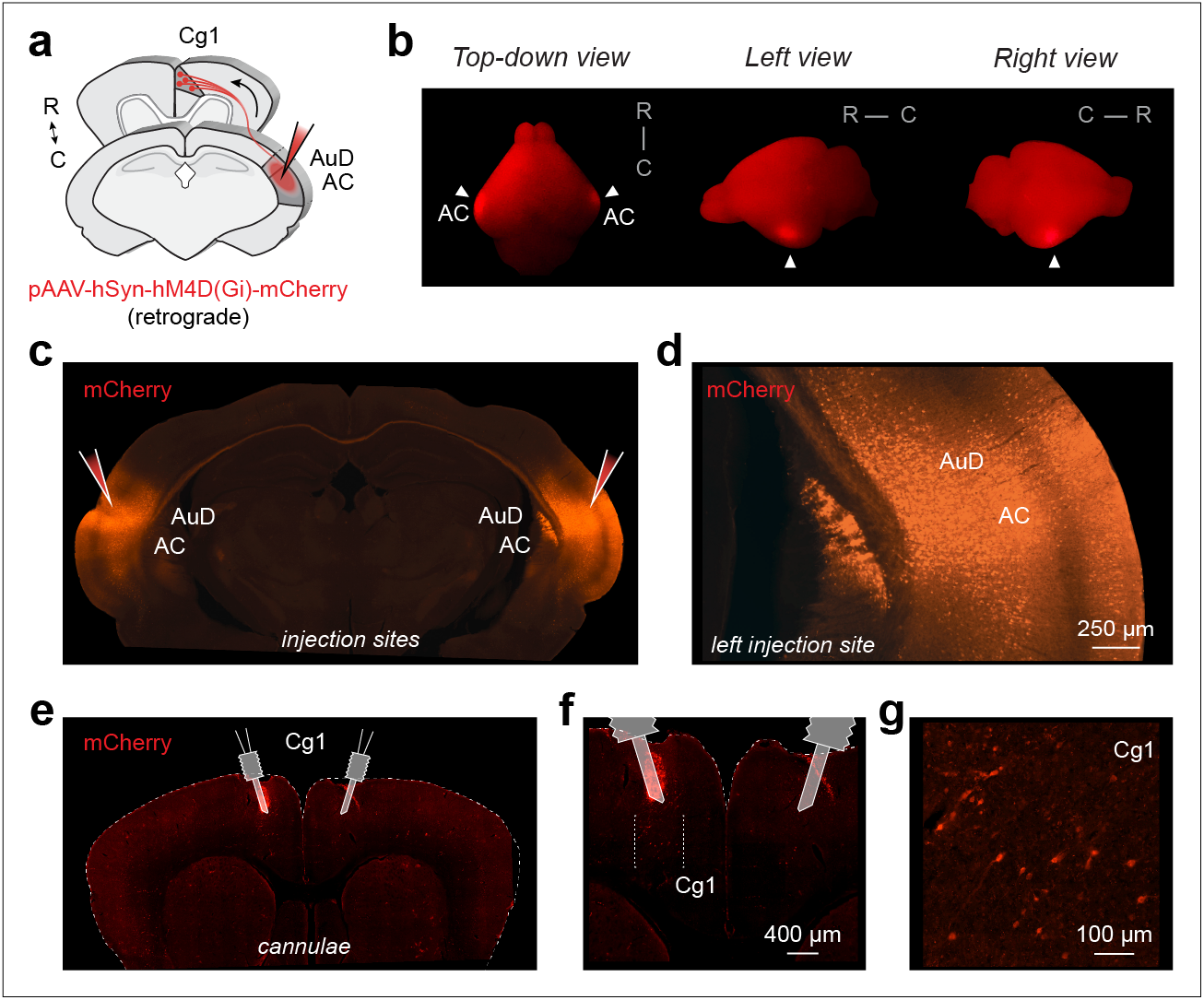
Confirmation of target locations for DREADDs inactivation experiment. (**a**) Schematic of DREADDs approach. To target cingulate cortex (Cg1) inputs to auditory cortex (AC, AuD), bilateral injections of a retrograde pAAV-hSyn-hM4D(Gi)-mCherry are made into auditory cortex (AC and AuD). (**b**) Whole-brain view (top-down, left-side, right-side) of example animal used in DREADDs experiment. White arrows indicate injection sites in auditory cortex (AC). (**c**) Brain slice showing bilateral injections of pAAV-hSyn-hM4D(Gi)- mCherry (retrograde) into AC. (**d**) High magnification of the left injection site in AC with labeled cell bodies. Scale bar represents 250 µm. (**e**) Brain slice showing site of bilateral cannulae implants above cingulate cortex (Cg1). (**f**) High magnification of targeted Cg1 region. (**g**) High magnification of expanded inset (white square in panel f) confirms Cg1 cell body labeling from injection sites in (c, d). Scale bar represents 100 µm.

**Supplemental Figure 6.**
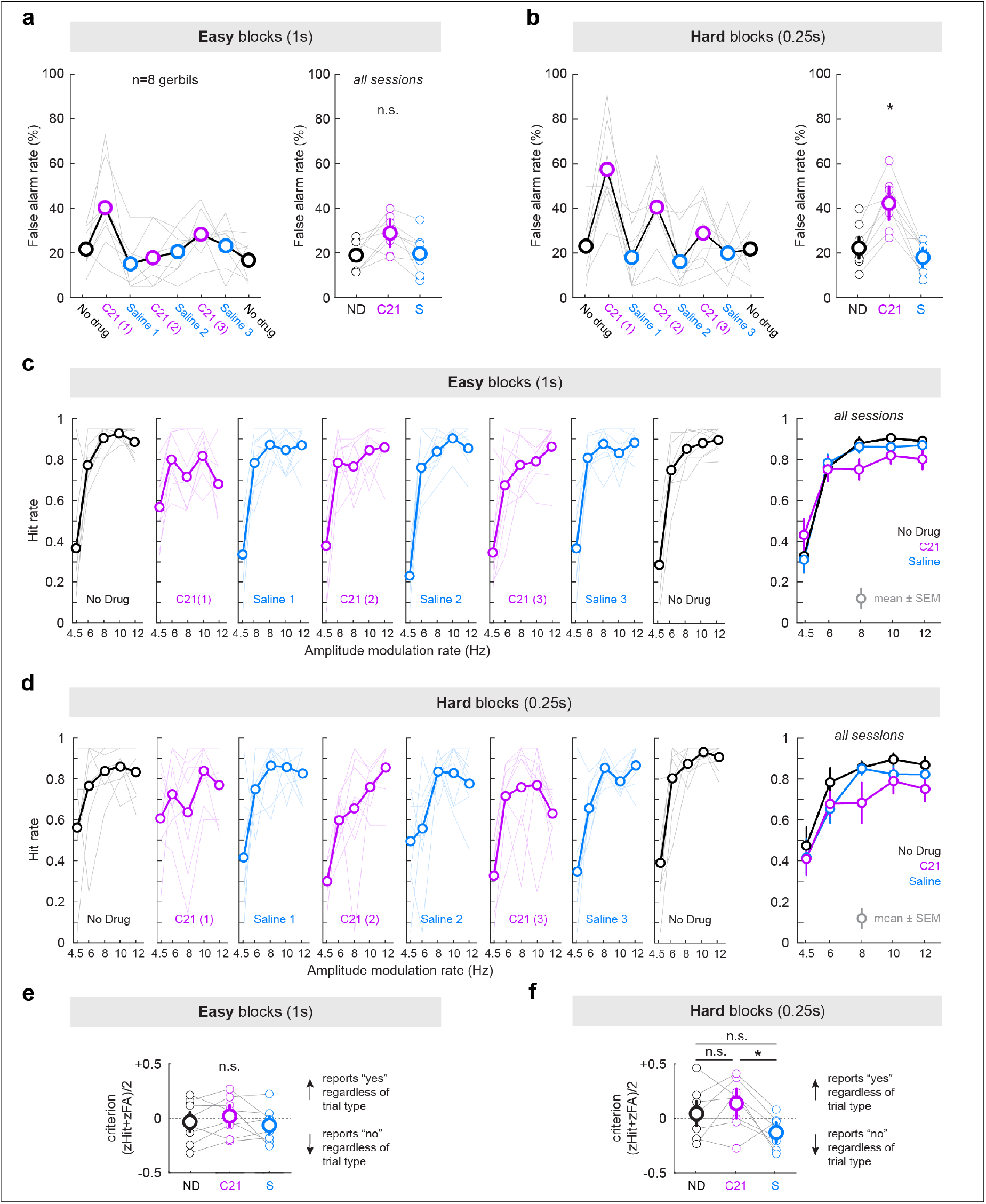
Inactivating Cingulate-to-Auditory cortex projections shifts behavioral response types for Hard blocks. (**a**) False alarm rate (%) across all testing days for Easy blocks (left panel) and all Easy block sessions combined (right panel). Large circles indicate the group average and grey lines indicate individual data. False alarms across infusion conditions for Easy blocks are not statistically different from one another (mixed model ANOVA followed by post hoc test for multiple comparisons: p=0.25-0.3). (**b**) False alarm rate (%) across all testing days for Hard blocks (left panel) and all Hard block sessions combined (right panel). False alarms across conditions for Hard blocks are significantly higher following C21 infusions compared to saline control (mixed model ANOVA: p<0.0001; post hoc test for multiple comparisons: p≤0.0006). (**c**) Hit rate across AM rate (Hz) for each individual infusion session for Easy blocks. The average Hit rate across AM rate for each condition is shown on the far right panel. Inactivating Cg-AC projections does not significantly alter Hit rates for Easy blocks (mixed model ANOVA followed by a post hoc test for multiple comparisons: p=0.7-0.9). (**d**) Hit rate across AM rate (Hz) for each individual infusion session for Hard blocks. The average Hit rate across AM rate for each condition is shown on the far right panel. Inactivating Cg-AC projections yielded a significant effect for Hard blocks compared to No Drug Hit rates (p=0.0007), though not compared to Saline Hit rates (p=0.3). (**e**) The average criterion ((zHit+zFA)/2) for each infusion condition for Easy blocks. Positive values indicate a more liberal response strategy (reports “yes” regardless of trial type) and negative values indicate a more conservative response strategy (reports “no” regardless of trial type). Inactivating Cg-AC projections did not shift performance strategy for Easy blocks (mixed model ANOVA followed by post hoc test for multiple comparisons: p=0.92-0.99). (**f**) The average criterion ((zHit+zFA)/2) for each infusion condition for Hard blocks. Inactivating Cg-AC projections did not shift performance strategy between C21 and No Drug Hard conditions (mixed model ANOVA followed by post hoc test for multiple comparisons: p=0.91), but did between C21 and Saline Hard conditions (p=0.02).

**Supplemental Figure 7.**
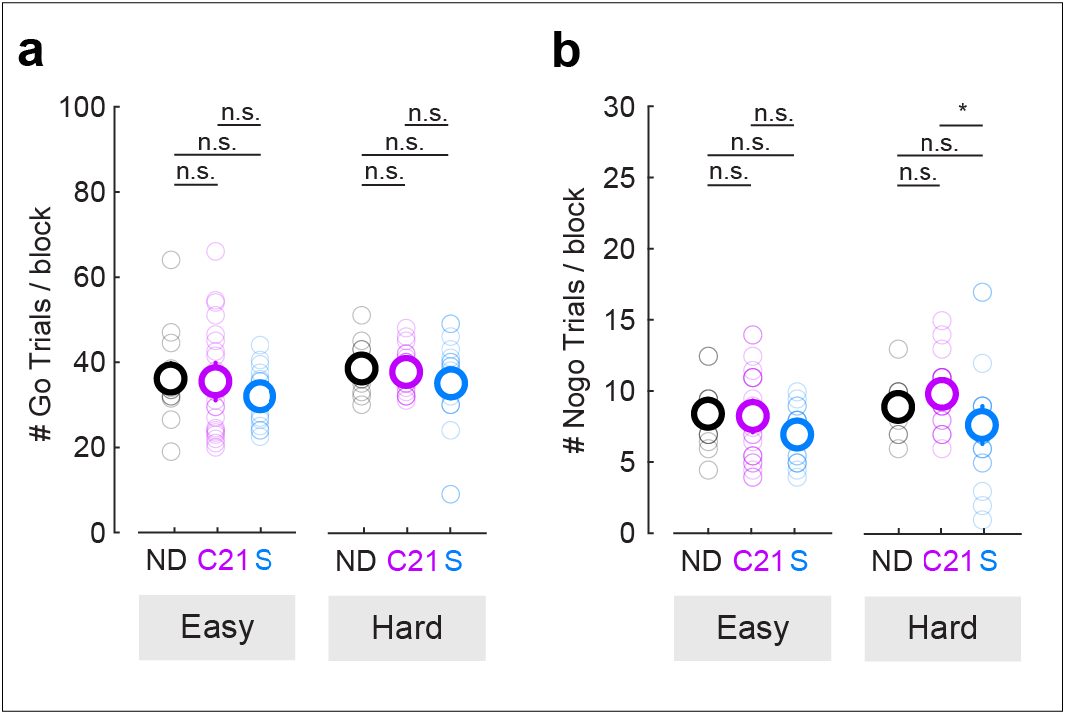
Performance deficits following inactivation of cingulate-to-auditory cortex projections (DREADDs) are not due to differences in trial number. (a) Number of Go trials for each block (Easy, Hard) for each infusion condition (No Drug, ND; Compound 21, C21; Saline, S). A mixed model ANOVA reveals no significant differences between groups (p=0.17). (b) Number of Nogo trials for each block (Easy, Hard) for each infusion condition (No Drug, ND; Compound 21, C21; Saline, S). Mixed model ANOVA followed by post hoc analysis for multiple comparisons reveal no significant differences between groups for Easy blocks (p=0.54-1) and between No Drug and C21 conditions (p=0.87) for Hard blocks (with the exception of Saline and C21: p=0.05).

**Supplemental Figure 8.**
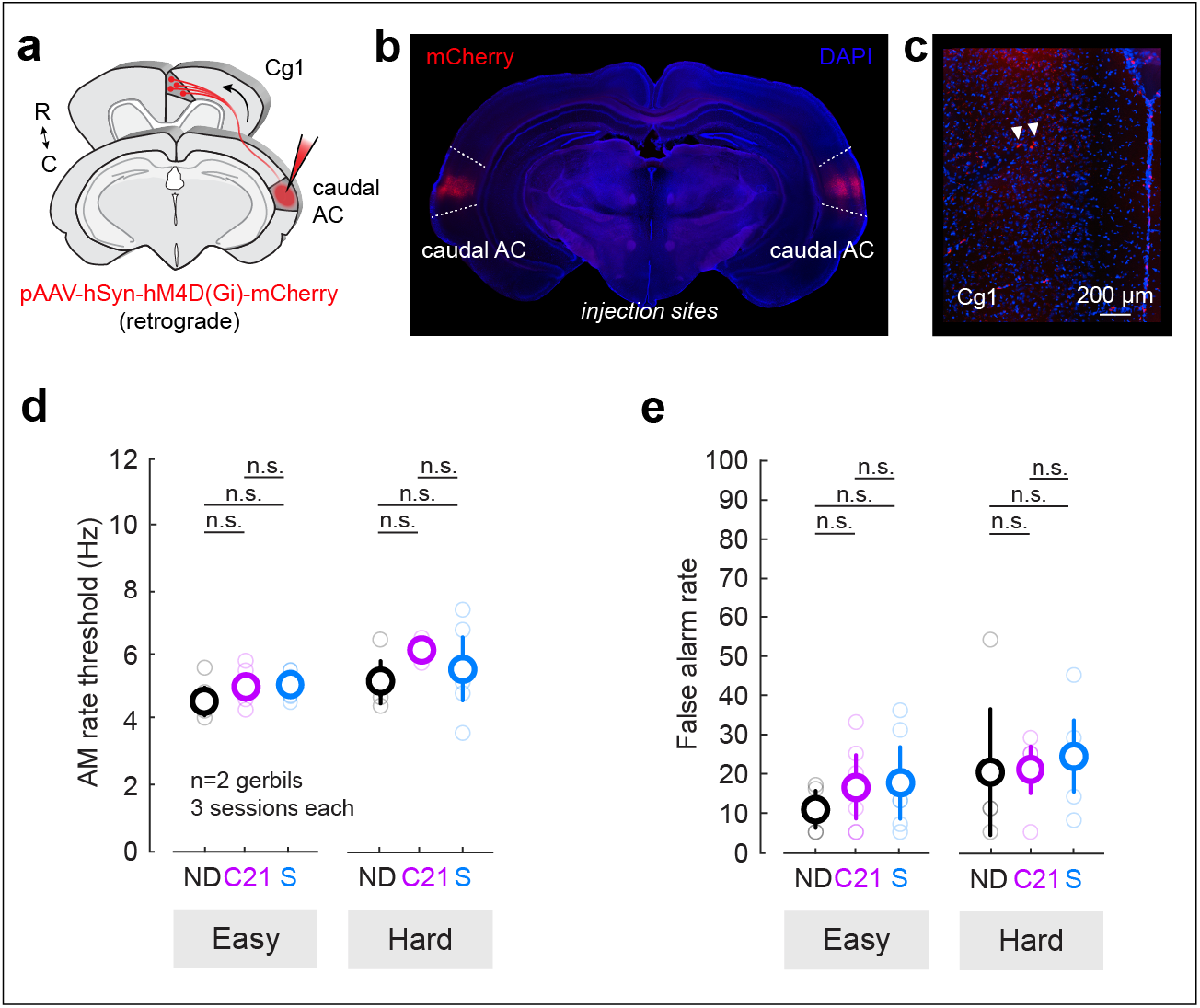
Inactivating cingulate projections to caudal auditory cortex does not alter perceptual performance on the Auditory effort task. (**a**) Schematic of DREADDs approach. Bilateral injections of a retrograde pAAV-hSyn-hM4D(Gi)-mCherry are made into caudal auditory cortex (AC; +1.4mm re: lambda). (**b**) Brain slice showing bilateral injections of pAAV-hSyn-hM4D(Gi)-mCherry (retrograde) into caudal AC. (**c**) Cross-section through Cg1 region. Sparse labeling of Cg1 cell bodies are present (white arrows). Scale bar represents 200 µm. (**d**) Average AM rate threshold (± SE) for all animals (n=2) and testing sessions combined (3 sessions per infusion condition). Thresholds remain unaffected following C21 infusions in Cg1 compared to No Drug and Saline controls for Easy or Hard blocks (mixed model ANOVA followed by post hoc test for multiple comparisons: p=0.4-1.0. (**e**) Average False alarm rates (± SE) for all animals (n=2) and testing sessions combined (3 sessions per infusion condition). False alarm rates remain unaffected following C21 infusions compared to No Drug and Saline controls for Easy or Hard blocks (mixed model ANOVA, p=0.68).

## Notes

### Competing Interest Statement

The authors have declared no competing interest.

